# *clipplotr* - a comparative visualisation and analysis tool for CLIP data

**DOI:** 10.1101/2021.09.10.459763

**Authors:** Anob M. Chakrabarti, Charlotte Capitanchik, Jernej Ule, Nicholas M. Luscombe

## Abstract

CLIP technologies are now widely used to study RNA-protein interactions and many datasets are now publicly available. An important first step in CLIP data exploration is the visual inspection and assessment of processed genomic data on selected genes or regions and performing comparisons: either across conditions within a particular project, or incorporating publicly available data. However, the output files produced by data processing pipelines or preprocessed files available to download from data repositories are often not suitable for direct comparison and usually need further processing. Furthermore, to derive biological insight it is usually necessary to visualise CLIP signal alongside other data such as annotations, or orthogonal functional genomic data (e.g. RNA-seq). We have developed a simple, but powerful, command-line tool: *clipplotr*, which facilitates these visual comparative and integrative analyses with normalisation and smoothing options for CLIP data and the ability to show these alongside reference annotation tracks and functional genomic data. These data can be supplied as input to *clipplotr* in a range of file formats, which will output a publication quality figure. It is written in R and can both run on a laptop computer independently, or be integrated into computational workflows on a high-performance cluster. Releases, source code and documentation are freely available at: https://github.com/ulelab/clipplotr.

## Introduction

Over the last twenty years, the study of RNA-protein interactions has been revolutionised by crosslinking and immunoprecipitation (CLIP) technologies (Lee and Ule 2018). There is now a wealth of publicly available CLIP data produced by multiple labs and also by consortia, such as ENCODE (Mukherjee et al. 2019; Van Nostrand et al. 2020). Databases have been established for the easy availability of processed CLIP data (Blin et al. 2015; Zhu et al. 2019) which can facilitate comparative exploratory analyses and integration of new experiments with the public corpus. Moreover, advances in the methodology have led to readily accessible protocols and wider uptake of CLIP experiments (Van Nostrand et al. 2017; Hafner et al. 2021; Porter et al. 2021; Buchbender et al. 2020; Lee et al. 2021). As a result, the questions now being addressed using CLIP experiments are often comparisons between different conditions, or between different RNA binding proteins (RBPs). A number of statistical approaches have been developed to assess differential binding using CLIP data, often based on extending differential gene expression methods originally designed for RNA-seq (Love et al. 2014; Liu et al. 2017; Wang et al. 2014; McIntyre et al. 2020). Alongside these bulk statistical analyses, it is crucial to visualise CLIP binding signals from multiple experiments in transcripts or regions of interest (ideally alongside orthogonal functional data such as RNA-seq, ribosome profiling or 3’-end sequencing) in order to support and understand differences in specific cases. Moreover, comparative analysis often involves iteration between visualisation and processing adjustments; most tools focus on one or the other, but *clipplotr* allows the user to do both when studying a region of interest.

Easy visualisation of CLIP data alongside orthogonal genomic data is crucial to guide biological interpretation of the RNA-protein interactions by contextualising binding sites or peaks with functional data. However, there are few options for experimentalists with limited bioinformatics experience to explore their data easily. SEQing is a tool that has recently been published to visualise iCLIP data and RNA-seq coverage (Lewinski et al. 2020). It is a locally-hosted, web-based tool that allows interactive exploration of the data tracks, similar to a genome browser and allows sharing of the session across group members. However, the CLIP and RNA-seq tabs can only be viewed in turn rather than simultaneously, preventing the user from easily identifying relationships at a glance. Moreover, SEQing only visualises the supplied data as is and does not perform any processing, hence making valid comparisons between datasets difficult and hindering the iterative nature of comparative analysis discussed earlier.

Crucially, there are two important considerations before processed CLIP data can be compared that preclude simply viewing the BED or BedGraph data tracks in a genome browser. First, the data from different experiments needs to be normalised to account for differences in library size using an approach appropriate to the study question. Second, the CLIP signal will generally benefit from being smoothed to aggregate crosslink data and highlight differences in binding patterns between experiments, conditions or RBPs. The need for this latter processing is generally underappreciated and can be a major problem for comparative CLIP visualisation.

To address these gaps and facilitate exploratory CLIP data analysis by the RNA community, we developed *clipplotr*: a self-contained command-line script that can be easily run with one command to produce publication-quality figures for defined genomic regions of interest. The tool simplifies visualisation of CLIP data alongside auxiliary and orthogonal data (e.g. RNA-seq, ribosome profiling or 3’-end sequencing) and transcript annotations from reference databases (e.g. Gencode or Ensembl). Most importantly, we have built in multiple normalisation and smoothing approaches for CLIP data that are essential to enable reliable comparisons.

## Results and discussion

Here we present the features of *clipplotr* in two use cases with a range of publicly available data from selected publications and the ENCODE Consortium (Zarnack et al. 2013; Van Nostrand et al. 2016, 2020). The latest CLIP technologies all identify nucleotide-resolution crosslink coordinates, which correspond to the position where the RBP crosslinked to the RNA. Depending on the method, this can then be identified through diagnostic events: either truncations (e.g. iCLIP, eCLIP) or mutations (e.g. PAR-CLIP) (Lee and Ule 2018; Chakrabarti et al. 2018). Processing of CLIP data is beyond the scope of *clipplotr* but is described in detail elsewhere (Chakrabarti et al. 2018; Busch et al. 2020) and can be performed by various computational pipelines available to run on local computing clusters (e.g. iCLIP: https://github.com/tomazc/iCount, eCLIP: https://github.com/YeoLab/eclip, PAR-CLIP: https://github.com/ohlerlab/PARpipe, nf-core: https://github.com/nfcore/clipseq (Ewels et al. 2020) or on webservers (e.g. iMaps: https://imaps.genialis.com). Here, all CLIP datasets were downloaded already processed to highlight the expected use of the tool.

### *clipplotr* generates a comprehensive and customisable visualisation with a single command

hnRNP C and U2AF2 (previously termed U2AF65) have been shown to compete directly to protect the transcriptome from the exonisation of *Alu* elements (Zarnack et al. 2013). The authors used iCLIP experiments to show that hnRNP C bound to cryptic splice sites suppressed the exonisation of *Alu* elements. However, loss of hnRNP C through siRNA knockdown experiments resulted in the expression of these *Alu* exons, demonstrated through RNA-seq experiments. Complementary iCLIP experiments demonstrated that U2AF2, a splicing factor, did not bind at the hnRNP C binding sites when hnRNP C was present, but upon knockdown of hnRNP C, there was increased U2AF2 binding at precisely these sites. This led to the model of direct competition between the two RBPs controlling *Alu* exonisation.

All of these findings are showcased in Fig. 1 with the unaltered output produced by *clipplotr*. We reproduce the example on the CD55 gene presented in the original paper, using all four of the tracks able to be generated by *clipplotr*. Input data are iCLIP BedGraph files for the crosslink track; RepeatMasker Alu BED files for the auxiliary track; RNA-seq coverage bigWig files for the coverage track; and a Gencode annotation GTF file for the annotation track. All the features of the plot annotated in Fig. 1 can be customised as desired using optional additional parameters. In the first panel - **the crosslink track** - the iCLIP BedGraphs have been normalised by library size to give a crosslinks-per-million calculation to permit valid comparisons. The signal has been smoothed with a rolling mean using a window size of 50 nt to show excellent concordance between replicates and reveal differences in crosslink signal, which represent binding regions, across the RBPs and conditions. Sets of experiments have been grouped together and coloured accordingly: duplicates of hnRNP C, U2AF2, and U2AF2 with hnRNP C knocked-down. The coordinates of the grey box can be specified and here is used to highlight sites of interest: in this case the competitive binding site. U2AF2 only binds to this site in the absence of hnRNP C, but in this context, it binds as strongly as hnRNP C did.

**Figure 1:**
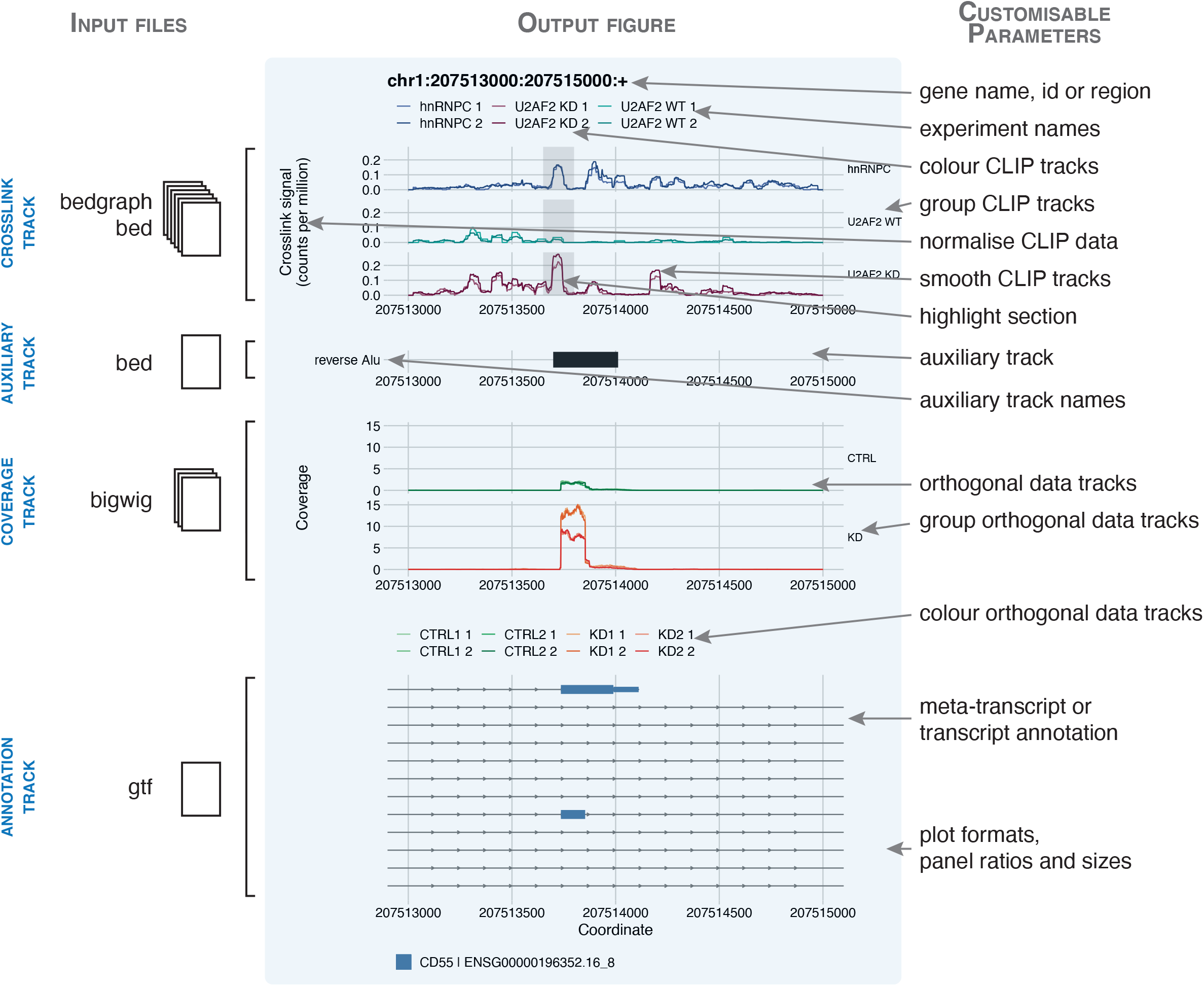
clipplotr outputs high-quality, easily customisable figures. A figure generated by a clipplotr using data from (Zarnack et al. 2013) is inset in blue, showing all four types of track that can be generated. The input file formats required for each track type is indicated to the left. On the right are annotated the customisable parameters that can be specified in the single clipplotr command.

The second panel - **the auxiliary track** - shows the location of inverted or reverse *Alu* elements (obtained from the UCSC table browser). The binding site where hnRNP C and U2AF2 compete is located at the 5’ end of the reverse *Alu* element - directly over the splice site.

The third panel - **the coverage track** - can be used to plot orthogonal genomic data. Here we show the coverage of RNA-seq data from three sets of experiments: four replicates of wild-type (CTRL) and two replicates each of two different knockdown siRNAs (KD1 and KD2). Again these have been grouped and coloured accordingly. There is a marked increase in expression of the *Alu* element in both knockdown conditions, indicating that the repression in the wild-type state has been relieved and the *Alu* element has been exonised.

The fourth panel - **the annotation track** - shows all the transcripts in the region of interest from the Gencode 34 annotation (from 2020). There are two transcripts that contain an exon matching the *Alu* element in the auxiliary track and the region of RNA-seq expression in the coverage track, but for the majority of CD55 transcripts this is an intronic region. The plot, inset in blue, is produced using one command, with aesthetics such as labels, colours, groupings, panel ratios and overall plot size all optionally customisable.

### clipplotr’s smoothing function highlights relevant changes in binding profiles

The importance of smoothing is demonstrated in Fig. 2 with the same hnRNP C and U2AF2 example as Fig. 1. Visualising the agreement of the two replicates in each group is not possible when viewing raw crosslinks as a bar graph, because the bars overlap rendering differences indistinguishable (Fig. 2A). In grey, we highlight the region containing the binding site where hnRNP C competes with and displaces U2AF2 in WT cells. In the raw data, the peak of crosslinking in this region is apparent in all conditions, but quantitative differences between conditions are less apparent because the signal from adjacent clustered crosslink sites are not aggregated. Instead, in U2AF2 after hnRNP C knockdown, other isolated positions downstream of this region with high signal also draw the eye (red arrowheads). Only after smoothing using a rolling mean with a 50 nt window (Fig. 2B) does the quantitative difference in the amount of crosslinking across the peak regions become clear: making it apparent that it is the amount of binding, but not the position of the binding, that changes. Although there are also increases in binding in the two downstream regions, it thus becomes evident that the primary site of competition between U2AF2 and hnRNP C is located in the region between 207,513,650 and 207,513,800 (the grey highlight box).

**Figure 2:**
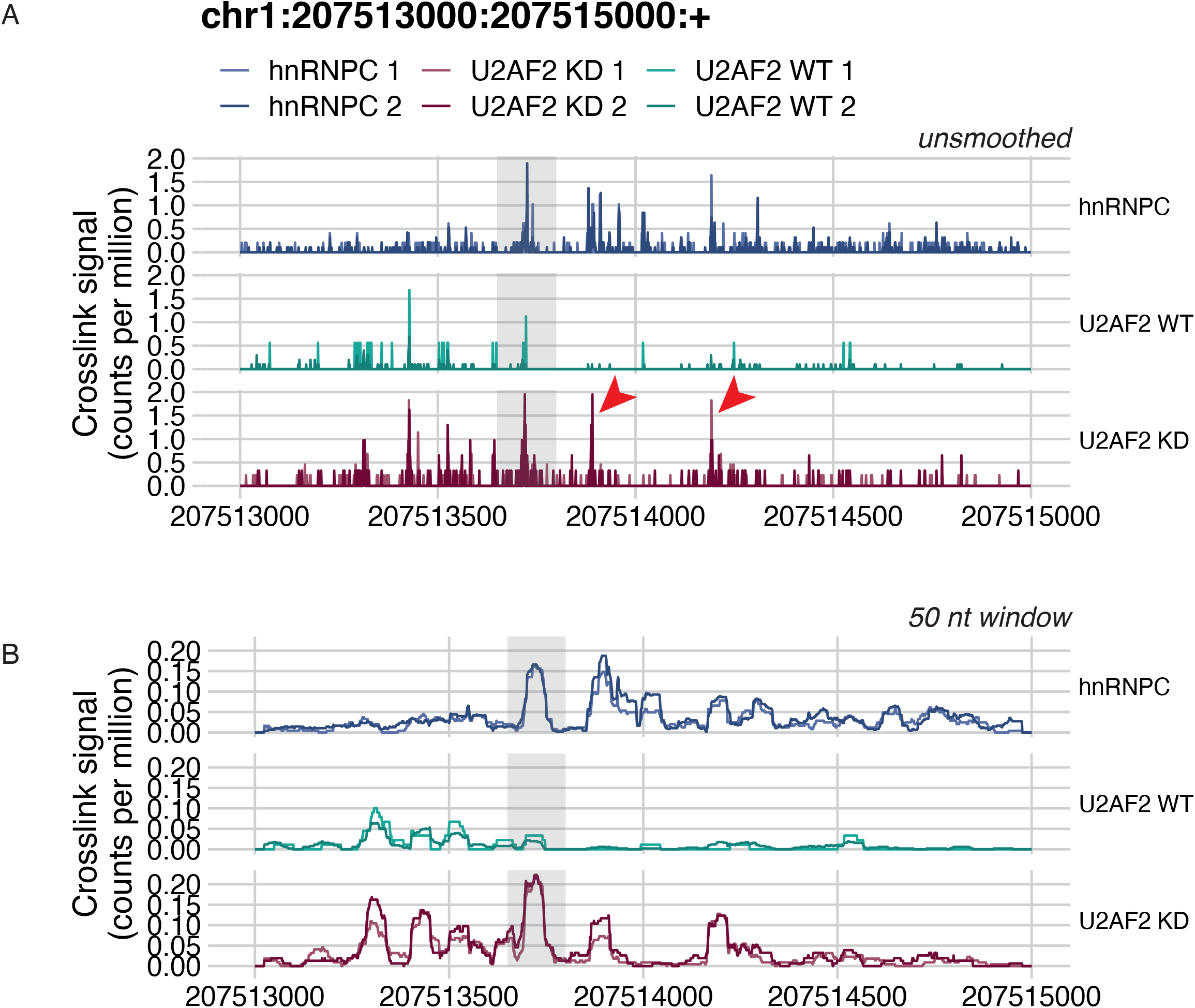
clipplotr’s smoothing functions highlight relevant changes in binding profiles. The same data is used as in Fig. 1. Highlighted in the grey box is the region of competition between hnRNPC and U2AF2 binding. (A) No smoothing has been applied using `--smoothing none`. The red arrowheads indicate downstream binding peaks that appear similar to that in the highlight box. (B) The signal has been smoothed using a rolling mean with a 50 nt window using `--smoothing rollmean --smoothing_window 50`.

Current CLIP experiments produce data that identify crosslink sites at single nucleotide resolution. However, when visualising these data over broad regions, such as whole genes, exons, or introns, smoothing is necessary to aggregate quantitative information from adjacent crosslink sites; otherwise the signal from individual crosslinks can become imperceptible, especially when comparing datasets. Furthermore, it is possible to introduce technical variation due to the many steps involved in CLIP library preparation (for example, from the uridine bias of UV crosslinking, from variation in RNase concentration or in cutting from the gel) that may result in the data not being fully reproducible at single nucleotide resolution, but become more so after aggregating or smoothing. We have included both a rolling mean (default) and a Gaussian approach as smoothing methods. The smoothing window size will depend both on the RBP and size of the region under study as well as the quality of the data and can be optimised to best aggregate the signal.

In this first use case we have shown how *clipplotr* can be used to normalise, smooth and compare binding of different RBPs in different conditions and, with the addition of annotation and orthogonal data, demonstrate their functional effects.

### *clipplotr’s* normalisation strategies allow exploration of multiple facets of the data

In the second use case (Fig. 3) we reproduce the finding of an RBFOX2 binding site on the NDEL1 transcript in the last intron of the gene, close to the 3’ UTR used as an example when the eCLIP method was first described (Van Nostrand et al. 2016). However, we use more recent eCLIP data in HepG2 and K562 cell lines produced by the same lab as part of the ENCODE Consortium (Van Nostrand et al. 2020). This allows us to showcase a possible use of the tool using exclusively publicly available processed data on sites of interest and how different datasets can be compared.

**Figure 3:**
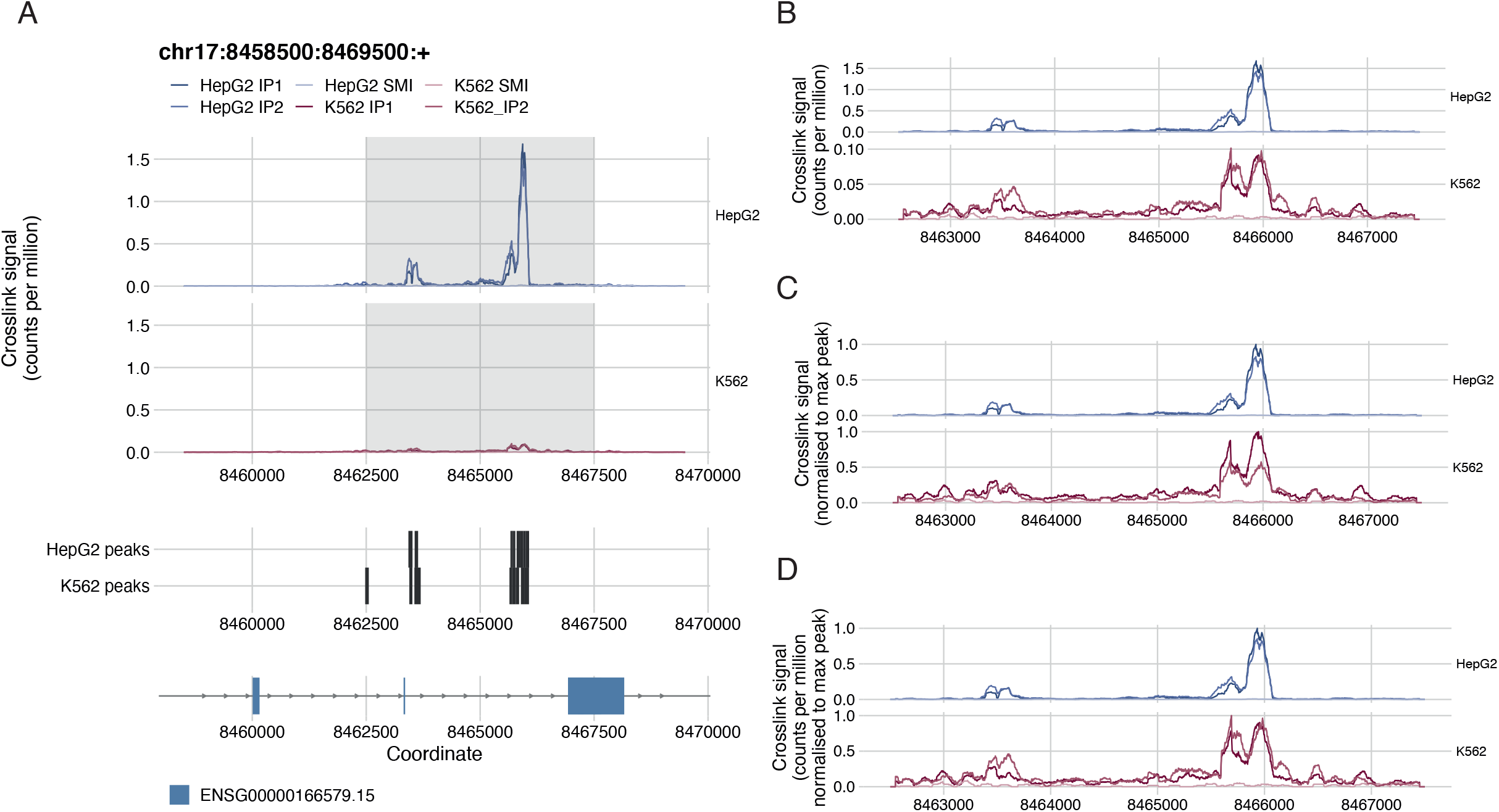
clipplotr’s normalisation functions allow exploration of multiple facets of data. (A) The figure generated by clipplotr using data from the ENCODE project showing the region from (Van Nostrand et al. 2016) with the CLIP data normalised by library size using `--normalisation libsize` (default). The region zoomed in in the subsequent panels is highlighted (B) The default library size normalisation is again used, but additionally the y-axis is scaled independently for each group using `--scale_y`. (C) The signal is normalised to the maximum peak in the region for each group using `--normalisation maxpeak`. (D) The signal is normalised first to library size and then by maximum peak in the region for each group using `--normalisation libsize_maxpeak`.

We use the CLIP, auxiliary and annotation track options from *clipplotr* to generate this image (Fig. 3A). Here, to complement the first use case, we show a “meta-transcript” annotation for the NDEL1 gene, which collapses all the exons across the transcripts to simplify visualisation. (Note that the coordinates differ slightly from the original figure as the newer ENCODE eCLIP data uses the hg38 sequence assembly, rather than hg19.) First, we have grouped the CLIP tracks by cell line and normalised by library size (Fig. 3A). Normalising by library size calculates a crosslinks-per-million value for each position, and accounts for the different sequencing depths of different experiments. It is immediately evident that there is a much stronger RBFOX2 binding signal in the HepG2 cell line compared to K562. This may reflect either differing expression levels of the NDEL1 transcript or technical variability of the two sets of experiments. However, the binding is still present, and moreover there is little binding seen at these sites in the size-matched input samples. Consequently, peaks have been called at similar sites for both cell lines as can be seen in the auxiliary tracks.

To visualise and explore the patterns of binding in more detail, we have included multiple options to normalise and scale the data: (i) do not normalise; (ii) normalise by library size (default); (iii) normalise to the maximum peak in each group; (iv) normalise by library size and scale the y-axis for each group; (v) normalise by library size and then to the maximum peak in each group; and (vi) apply a custom normalisation factor. The y-axis label is automatically adjusted based on the method. The importance of normalising by library size prior to comparing different experiments is well known, but we have kept the option not to as it may be useful to examine the raw signal in single experiments. In Fig. 3B, we show the effect of (iv): so that the signal for each group fills its respective plot (Fig. 3B). This preserves the information that the two sets of experiments are an order of magnitude different in the strength of the signal, but also allows delineation of the peak morphology. Another approach often used is (iii): this should allow a more easy comparison of the relative differences between the two groups of experiments as the crosslink signal is scaled from 0 to 1 (Fig. 3C). However, this approach should be used with caution: for K562, it appears as though replicate 1 has half the signal of replicate 2, whereas in Fig. 3B we can see that the two have comparable signal when normalising by library size. Thus the disparity is accounted for by different library sizes: 10,942,658 v 20,298,696 reads. When examining much larger regions, there may be enough signal across the region to account for library size differences when using this approach, but often for the more targeted analysis undertaken for CLIP data it can be confounded. If it is important to show relative differences in this way, we advocate using (v) for CLIP data as shown in Fig 3D. Here the profile is identical to Fig. 3B, but the y-axis values now allow easy quantification of relative differences between groups.

In this second use case, we have highlighted how the different *clipplotr* normalisation strategies can each be used to derive different, complementary insights into the data and can be selected based on the question or the specific effects of RBP binding under study. We have also shown a potential pitfall of normalisation to the maximum peak and recommend an alternative option.

## Conclusions

Visual inspection of sequencing data is an important part of the data analysis process. When such data are produced to provide nucleotide-resolution information, such as is the case for iCLIP, it can be challenging to visualise its quantitative aspects at the level of genes or other broader genomic regions. To solve this challenge, we have developed *clipplotr*, a command-line tool to facilitate comparative visual exploration of CLIP and orthogonal datasets. We provide a range of normalisation, smoothing and visualisation options to ensure appropriate comparisons can be easily undertaken. It is straightforward to customise all these options, and the many aspects of visual presentation, through the command line parameters. Equally, sensible default options have been established so that the tool will run with minimal user input. We have already found the tool very valuable also for visualisation of mNET-seq data, and we believe it will be of use to visualisation of many other high-resolution data apart of CLIP, such as for example studies of RNA structure (SHAPE-map, etc.), protein-DNA interactions (ChIP-exo), polymerase and ribosome-binding (NET-seq, Ribo-seq), and similar. This simple-to-use tool we hope will thus enable seamless data analysis, while also leading to plots that are of a standard to be included in a published figure.

## Materials and Methods

### Implementation and installation

*clipplotr* is publicly available under an MIT licence and maintained on GitHub (https://github.com/ulelab/clipplotr), where there is also comprehensive documentation. It is implemented in R (v. 4.0.2) using the R packages optparse, data.table, ggplot2, ggthemes, cowplot, patchwork, zoo and smoother, and the Bioconductor packages rtracklayer and GenomicFeatures. A Conda environment YAML is provided to generate a virtual environment which will install R and all the necessary dependencies. Alternatively, if the user already has R installed on their system, a helper R script is provided to install these packages. A small test dataset and command is also available to confirm correct installation by generating the plot shown in Fig. 1. We have tested *clipplotr* on macOS, Linux and Windows systems.

### Usage

All arguments to *clipplotr* can be passed at the command line. In addition to the usage documentation, details of all the clipplotr parameters, possible arguments and defaults can be accessed using `clipplotr --help`. The minimum input requirements for *clipplotr* are: (i) a set of CLIP crosslink position tracks in BED or BedGraph format (using `--xlinks`); (ii) a GTF file with the reference annotation, e.g. from GENCODE (using `--gtf`); (iii) a gene or genomic region of interest (using `--region`); and (iv) a filename for the output plot (using `--output`). This will produce a minimal plot that contains the crosslink and annotation tracks.

From the majority of analysis pipelines, either BED or BedGraph format files are usually produced in which each entry in the file is a genomic position and the score the number of crosslinks detected at that position. As BedGraph files do not contain strand information, it is common with iCLIP data for this to be encoded within the score: a positive value indicating the crosslink is on the positive strand and a negative value indicating it is on the negative strand (König et al. 2010). Preprocessed publicly-available crosslink data is also usually available in one of these file types (Van Nostrand et al. 2020; Zhu et al. 2019; Blin et al. 2015). Experiment names, colours and groupings can all be specified by the user, or automatically generated from the filenames. The reference annotation GTF file can be obtained from commonly used resources such as the GENCODE project (Frankish et al. 2019) or Ensembl. The first time *clipplotr* is run, it will generate and save an SQL database from the provided annotation file and use this to speed up future runs using the same annotation. The annotation tracks can either be plotted at the “gene” or transcript level. At the “gene” level, a single meta-transcript is plotted for each gene: this contains all annotated exons of the gene across all transcript isoforms. At the transcript level (default), all annotated transcript isoforms in the region are plotted and coloured by the gene with which they are associated. The region of interest can either be specified either by gene name, gene id, or using genomic coordinates. This will also be used as the title of the plot. The output plot can be generated as either a PDF or PNG file; the format is determined from the supplied filename extension.

Optionally, auxiliary tracks and coverage tracks can also be plotted for the same genomic region to relate CLIP signal to other complementary data as shown in Fig. 1 and Fig. 3. Auxiliary tracks are supplied as BED files and can for example be used to show either relevant genomic features (e.g. reverse *Alu* elements, Fig. 1), or CLIP features (e.g. peaks called using the CLIP crosslinks plotted in the crosslink tracks, Fig. 3). If a BED9 format file is supplied then the interval is coloured accordingly. Coverage tracks can be supplied as BigWig files and are plotted without further processing. As for the CLIP tracks, names, colours and groupings can all be specified by the user. If these optional tracks are included, they are dynamically scaled so all tracks appropriately fit the plot page size, however, the user can also specify precise ratios for the four tracks and the page size to their exact requirements..

As an full example, the detailed plot shown in Fig. 1 was generated using the “one-line” command:

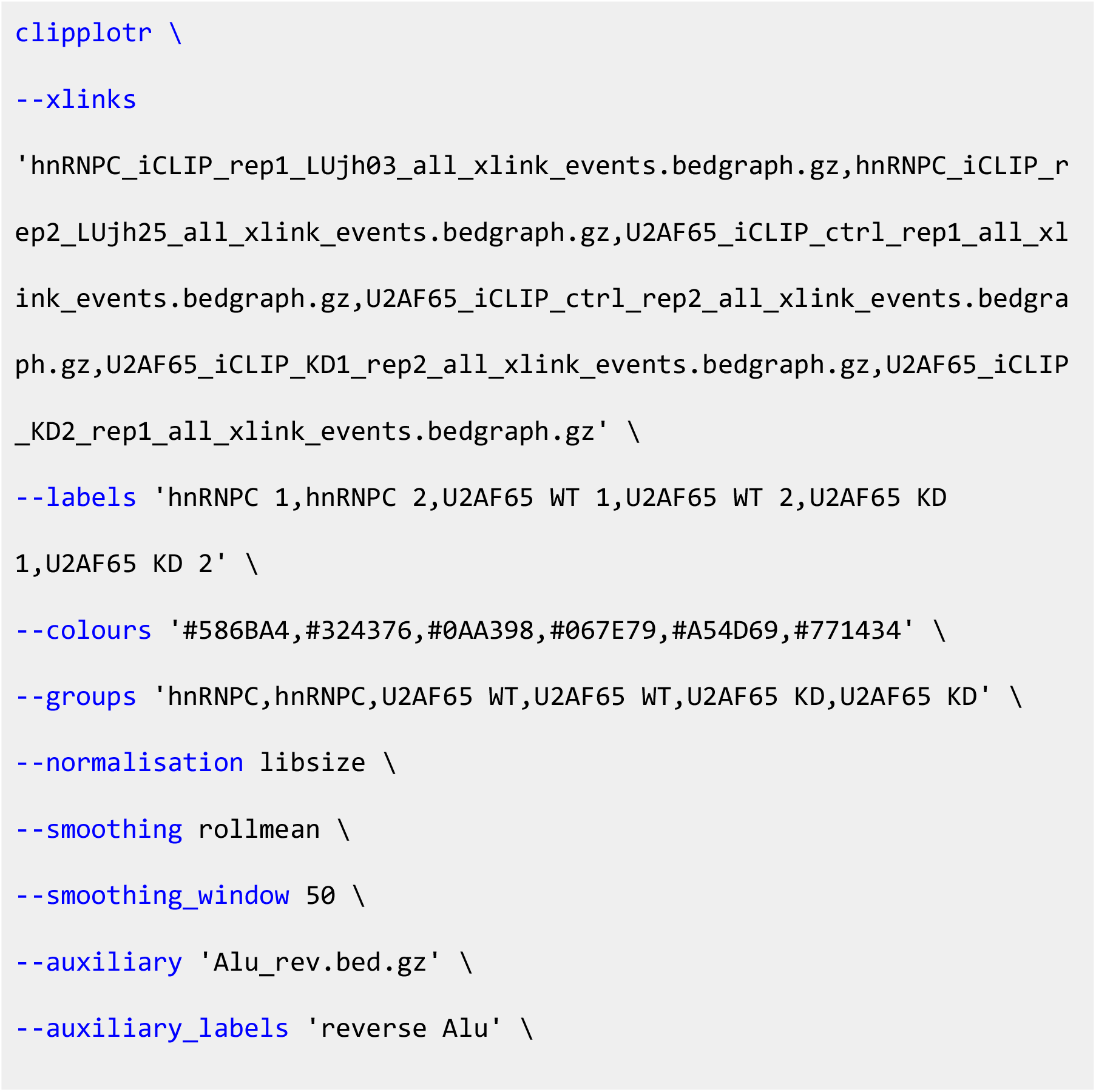

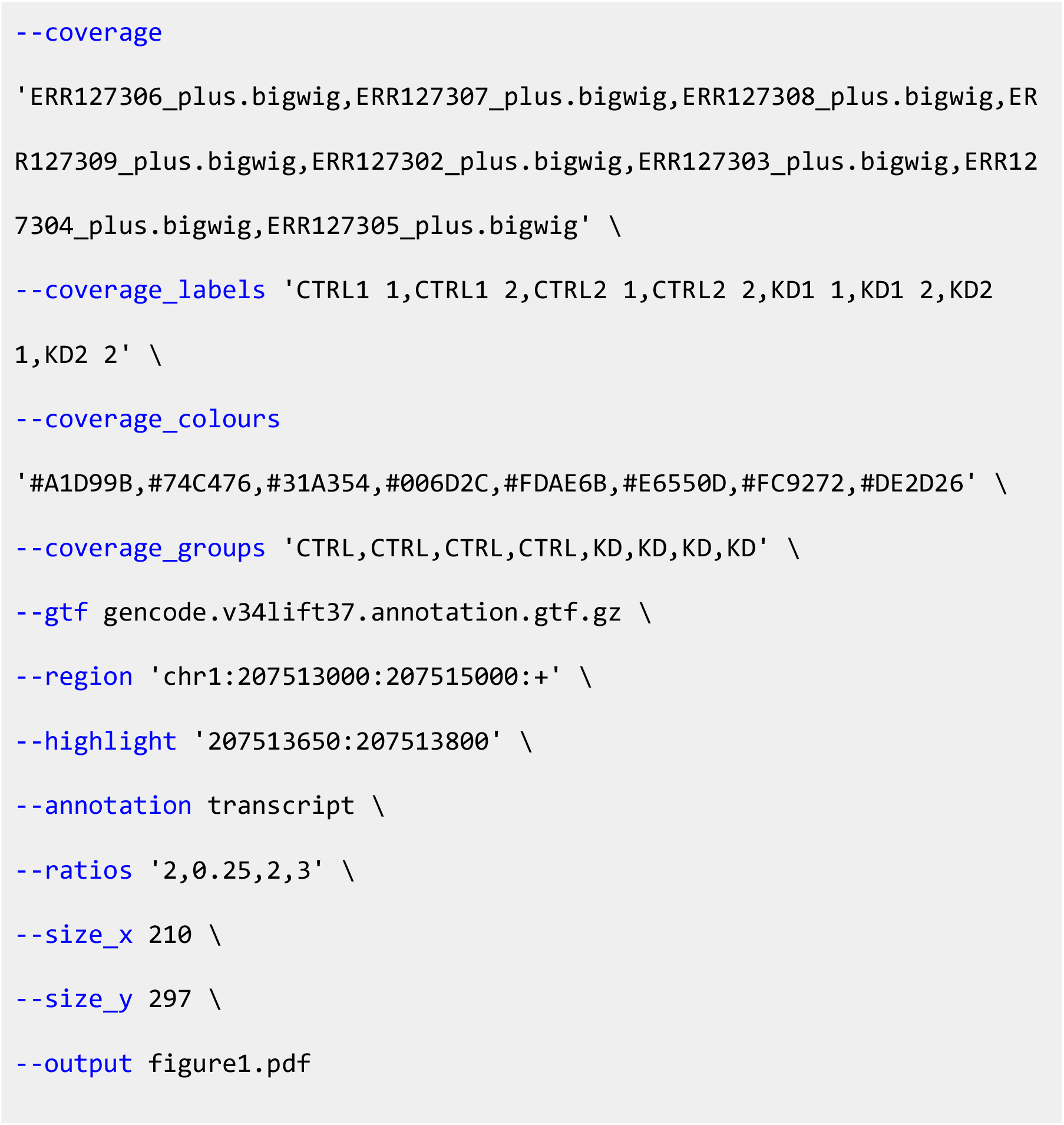

The long forms of the parameters are shown here for ease of comprehension, but short forms can also be used for the majority (e.g. `-x` or `--xlinks`). As described earlier, not all parameters need arguments to be provided at the command line as we have implemented sensible default options: the minimum requirements are `--xlinks`, `--gtf`, `--region` and `--output`.

### Data processing for use cases

We deliberately chose to use processed data wherever possible to focus on the main use case of *clipplotr*. In the first use case, we reproduced Fig. 1C from (Zarnack et al. 2013). All iCLIP crosslink BedGraph tracks were downloaded as processed data from ArrayExpress E-MTAB-1371. The *Alu* BED file was downloaded using UCSC Table Browser from the RepeatMasker track. The strands were swapped to make a reverse *Alu* BED file. Processed RNA-seq data were not available from ArrayExpress E-MTAB-1147, so were downloaded as raw FASTQ files (accession numbers ERR127302-9). Reads were trimmed using TrimGalore v. 0.6.4_dev (https://github.com/FelixKrueger/TrimGalore) and mapped to the GRCh37 genome assembly with the Gencode v34-lift37 GTF annotation using STAR v. 2.7.4a (Dobin et al. 2013). BIGWIG coverage tracks normalised to reads-per-million were created using STAR and bedGraphToBigWig from UCSC tools. The annotation GTF was the same as used for RNA-seq mapping.

For the second use case, we reproduced the results shown Fig. 1D from (Van Nostrand et al. 2016), but using more recent eCLIP data in HepG2 and K562 cell lines from the ENCODE Consortium (Van Nostrand et al. 2020) to showcase comparisons in binding between experiments and cell lines. eCLIP data was downloaded as processed data from the ENCODE portal (https://www.encodeproject.org). For the HepG2 cell line, crosslink BED files were from accession numbers ENCFF239CML, ENCFF170YQV, ENCFF515BTB and the peak BED file from ENCFF871NYM; and for the K562 cell line, crosslink BED files from ENCFF537RYR, ENCFF296GDR, ENCFF212IIR and the peak BED file from ENCFF206RIM. The basic annotation GTF was downloaded from Gencode (v34).

A snakemake pipeline script for the RNA-seq processing and the *clipplotr* commands to generate these use case plots are available at https://github.com/ulelab/clipplotr/examples.

## Acknowledgements

We would like to thank members of the Ule and Luscombe Labs for testing the tool and providing user feedback during its development, in particular Andrea Elser, Martina Hallegger and Flora Lee. This research was funded in whole, or in part, by the Wellcome Trust (FC010110; 215593/Z/19/Z). For the purpose of Open Access, the author has applied a CC BY public copyright licence to any Author Accepted Manuscript version arising from this submission. This work was supported by the Francis Crick Institute which receives its core funding from Cancer Research UK (FC010110), the UK Medical Research Council (FC010110), and the Wellcome Trust (FC010110). AMC was supported by a Wellcome Trust PhD Training Fellowship for Clinicians Award (110292/Z/15/Z) and is currently supported by a Crick Postdoctoral Clinical Fellowship. This work was also supported by Wellcome Trust Joint Investigator Awards (215593/Z/19/Z) to JU and NML. NML is a Winton Group Leader in recognition of the Winton Charitable Foundation’s support towards the establishment of the Francis Crick Institute. NML is additionally supported by core funding from the Okinawa Institute of Science & Technology Graduate University.

